# Effect of cadmium accumulation on the performance of plants and of herbivores that cope differently with organic defences

**DOI:** 10.1101/403576

**Authors:** Diogo Prino Godinho, Helena Cristina Serrano, Anabela Silva, Cristina Branquinho, Sara Magalhães

## Abstract

Some plants can accumulate in their shoots metals that are toxic to most other organisms. This ability may serve as a defence against herbivores. Although both metal accumulation and the production of organic defences may be costly to the plant, the two mechanisms may interact on their effect on herbivores. However, this interplay between metal-based and ‘classical’ organic defences remains overlooked.

To fill this gap, we studied the interactions between tomato (*Solanum lycopersicum*), a plant that accumulates cadmium, and two spider-mites, *Tetranychus urticae* and *T. evansi* that respectively induce and suppress organic plant defences, measurable via the activity of trypsin inhibitors. We exposed plants to different concentrations of cadmium and measured their effect on mites and plants. The oviposition of both spider-mite species was higher on plants exposed to low concentrations of Cd than on control plants but decreased at concentrations above 0.5 mM. Therefore, herbivores with contrasting responses to organic defences responded similarly to metal accumulation by the plants. On the plant, despite clear evidence for Cd accumulation, we did not detect any effect of Cd on traits that reflect the general response of the plant, such as biomass, water content and carbon/nitrogen ratio. Still, we found an effect of Cd supply upon the quantity of soluble sugars and leaf reflectance changes that may reflect structural modifications in the cells. In turn, these changes in plant traits interfered with the performance of spider mites feeding on those plants.

Additionally, we show that the induction and suppression of plant defences by spider mites was not affected by Cd supply to the plants. Furthermore, the effect of metal supply on spider-mite performance was not affected by previous infestation. Together, our results suggest no interaction between metal-based and organic plant defences, on our system. This may be useful for plants living in heterogeneous environments, as they may use one or the other defence mechanism, depending on their relative performance in each environment. This may be relevant to studies on the interactions between herbivores and plants, from physiology to ecology.

## Introduction

Plants are exposed to an array of abiotic and biotic stresses. The mechanisms that allow them to survive these adversities impose physiological and structural transformations that can be costly to the plants affecting negatively their growth and fitness (Boyer 1982, Wang et al. 2003). One such stress is that of soils with high bioavailable metal concentrations, either naturally (geochemical anomalies) or due to anthropogenic activities. Although high bioavailable metal concentrations are toxic to most organisms, some plant species or populations, termed metallophytes, thrive in such environments. They achieve this either by limiting metal translocation to upper parts (excluders), or by storing metals in their above parts (accumulators; Baker 1987). However, these strategies entail costs that may be reflected in the plant performance, namely in plant biomass, on the water content of the aerial parts and/or on the root to shoot ratio (Kastori et al. 1992, Das et al. 1997, Larbi et al. 2002, Chaffei et al. 2004, Devi et al. 2007). In addition, the stress caused by metal toxicity may lead to disturbances in the carbon and nitrogen metabolisms affecting the nutritional status of various plant parts (Larbi et al. 2002, Chaffei et al. 2004, Wahid et al. 2007) and also changing the accumulation of soluble sugars in the plants, either leading to increased (Devi et al. 2007, Rosa et al. 2009, Mishra et al. 2014) or decreased (Scheirs et al. 2006, Shackira and Puthur 2017) concentrations in the shoots. These physiological changes in the plant may also affect the performance of herbivores infesting those plants (White 1984, Scheirs et al. 2006).

Besides being costly for the plant, metals are highly toxic to herbivores as well. Indeed, due to the elemental nature of metals, they cannot be degraded by chemical counter-defenses of the herbivores (Boyd 2004). Therefore, metal accumulation by the plants may be detrimental to herbivores (Martens and Boyd 1994, Boyd and Moar 1999, Behmer et al. 2005, Kazemi-Dinan et al. 2014) and can thus serve as a defence against herbivores (Boyd 2004, Poschenrieder et al. 2006, Horger et al. 2013). If this strategy is coupled with the production of organic defences, a possible joint effect would give accumulating plants an advantage over non-accumulating competitors (Boyd 2007). Indeed, metal exposure may directly increase the activity of some organic plant defences, such as proteases (Pena et al. 2006, Lin et al. 2010), having possible indirect effects on herbivores. However, because both types of defences may be costly to the plant, if metal-based defences are effective, the selection pressure upon the production of organic plant defences may be reduced. Indeed, some studies show that metal-accumulating plants produce fewer organic defences upon pathogen attack when they are supplied with metals (Farinati et al. 2011, Fones et al. 2013). This suggests a trade-off between metal-based and organic defences although more evidence is needed to establish causality and determine its prevalence, including its potential extension to herbivores.

Although most herbivores induce the production of organic plant defences (Karban and Myers 1989, Walling 2000, Awmack and Leather 2002), some are able to suppress them (Musser et al. 2002, Abramovitch et al. 2006, Sarmento et al. 2011). Likewise, metal defences vary in their effects upon herbivores. For example, the effectiveness of metal accumulation as an anti-herbivore defence varies with herbivore feeding guilds (Jhee et al. 2005, Vesk and Reichman 2009, Konopka et al. 2013), and between specialist and generalist herbivores (Kazemi-Dinan et al. 2014). However, it is yet unknown whether metal-based defences affect differently herbivores that induce or suppress organic defences. Addressing this issue would also provide insight into the potential interaction between metal-based and organic defences.

The system composed of tomato plants (*Solanum lycopersicum,* L.) and herbivorous mites is ideal to test the abovementioned issues. When growing on soils with cadmium (Cd), tomato plants show higher tolerance than other species (Bingham et al. 1973, Khan and Khan 1983, Kuboi 1986). Moreover, when supplied with this metal, tomato plants accumulate it on their shoots, sometimes over the hyperaccumulation threshold (Gratão et al. 2008, López-Millán et al. 2009). Among spider mites, *Tetranychus urticae* is negatively affected by metal accumulation of some host plants (Jhee et al. 2005, Quinn et al. 2010), but knowledge on other species remains elusive. Additionally, different species within the Tetranychidae show contrasting effects on the organic defences of the plant. Indeed, *T. urticae* induces the production of plant defences such as proteinase inhibitors, leading to lower performance of herbivores infesting tomato plants (Li et al. 2002, Ament et al. 2004, Kant et al. 2004). In contrast, *T. evansi* suppresses the production of such defences (Sarmento 2011, Alba 2015), leading to higher performances of herbivores on subsequent infestations (Sarmento et al. 2011, Godinho et al. 2016). These differences allow unravelling the possible interactions between organic and metal accumulation defences. To this aim, we assessed the effect of Cd accumulation on the performance of tomato plants and on the spider mites that infest those plants. Additionally, we evaluated the effect of herbivory on plant defences and subsequent infestations by spider mites on plants exposed to different Cd concentrations.

## Materials and methods

### Biological material and rearing conditions

#### Plants

Tomato plants (*Solanum lycopersicum*, var. Moneymaker) were sowed in a climate chamber (25 °C, photoperiod 16/8 h (light/darkness)), in a soil/vermiculite mixture (4:1) and watered 3 times per week for the first two weeks. In the 3^rd^ and 4^th^ weeks, plants were treated with a Cd solution, of a given concentration. To this aim, plants were watered once a week with tap water and twice a week with 60 mL of a Cd chlorine solution with two ranges of concentrations: a wide range 0, 0.01 mM, 0.1mM, 0.5 mM, 1 mM, 2 mM, or 10 mM; and a narrow range 0, 0.1mM, 0.25 mM, 0.5 mM, 0.75 mM, 1 mM, or 1.5 mM. Using the wide range, we tested the effect of high Cd concentrations on the plant and on spider mites. Using a narrow range allowed us to measure plant and spider mite traits with higher resolution.

#### Spider mites

*Tetranychus urticae* was collected from tomato plants in Portugal in 2010 and reared on bean plants (*Phaseolus vulgaris*, L.) since then (Clemente et al. 2016). In January 2016 a sub-set of the population (>300 mated females) was transferred to tomato plants and maintained on this host for 6 generations before being used in the subsequent experiments. *Tetranychus evansi* was collected from *Datura stramonium*, L. in 2013 in Portugal and reared on tomato plants ever since (Zélé et al. 2018). The two species were maintained separately in plastic boxes containing two entire tomato plants, in a climate chamber with conditions identical to those of the plant growing compartment (25 °C, photoperiod 16/8 h (light/darkness)). Once a week, one plant was removed, and its leaves were cut and placed on top of the leaves of a new plant, allowing spider mites to migrate to clean plants. To ensure that females used in the experiments were approximately of the same age, adult females where isolated on separate leaves and allowed to lay eggs for 48 h. 12 days later, the adult females resulting from these cohorts were used in the experiments.

### General methodology

The performance of plants and spider mites was assessed using plants exposed to both ranges of Cd supply. The same leaf (3^rd^ from below) was used for every assay in order to control for the effect of leaf age.

#### Plant performance

Because the plant material collected was not enough to use in every assay, measurements were performed with different plants: Plants exposed to the wide range of concentrations (0 to 10 mM, N=6 per Cd concentration) were used to determine Cd accumulation on the leaf, as well as the amount of calcium (Ca) and magnesium (Mg). From the narrow range (0 to 1.5 mM), half the plants (N= 6 per Cd concentration) were used to obtain the biomass parameters (root/shoot; specific leaf area and water content), and the remaining to measure the amount of soluble sugars and to determine the carbon (C) to nitrogen (N) ratio. However, for each plant, and before any destructive assay, we determined the spectral reflectance of the leaf, a non-invasive method that provides a general assessment of plant stress (Carter 1993, Carter and Knapp 2001).

##### Spectral analysis

The spectral reflectance was measured on one leaf from each plant, 5 measurements per leaf, using a UniSpec spectroradiometer (PP-Systems, Haver Hills, MA, USA). The spectral data originated by these measurements was analysed by calculating spectral reflectance factors (R) for each wavelength (between 300.4 and 1148.1 nm with intervals of 3.4 nm). These factors were obtained by normalizing the reflected radiation from the leaves by a reflectance white standard. Several vegetative indices can be determined using reflectance data and used as a proxy of plant stress, being the most commonly used the Normalized Difference Vegetation Index (NDVI) as it reflects the efficiency of the photosynthetic system (Sridhar et al. 2007). Therefore, we here measured NDVI ((R810-R680)/(R810+R680)). In addition, we measured the SC index, which is representative of structural changes in leaf cells caused by accumulation of Cd (R1110/R810; Sridhar et al. 2007). Moreover, as it has been proposed that plants share similar responses to UV-B light exposure and herbivory, such as phenolic compounds (Roberts and Paul 2006, Izaguirre et al. 2007), we also analysed the spectral data regarding those wavelengths. For that we averaged, for each plant, the spectral reflectance factors of all UV-B wavelengths (R300.4 – R313.9).

##### Cadmium, Calcium and Magnesium quantification

One leaf from each plant was dried for 72 hours at 60 °C until constant mass and uniformly grounded. The elements were then quantified using Inductively Coupled Plasma - Atomic Emission Spectrometry (ICP- AES), after nitric acid digestion, with a detection limit of 0.1 µg/L.

##### Root to Shoot ratio, Specific Leaf Area and Plant water content

All leaves and roots of each plant were collected, then the area of each leaf was measured with a laboratory leaf meter (Li-Cor Biosciences). Next, the fresh weight of leaves and roots was obtained. Each leaf and the roots were then separately dried for 72 hours at 60 °C until constant mass and again weighted. The ratio between the dry weight of the roots and the dry weight of the leaves (root/shoot) was determined as well as the specific leaf area (SLA, total leaf area / total leaf dry weight) and plant water content (fresh weight – dry weight / fresh weight).

##### Carbon to Nitrogen ratio

One leaf from each plant was dried at 60 °C until constant mass and again weighted. The total carbon (C) and nitrogen (N) contents (grams of C or N per 100g of leaf dry weight) of each leaf was determined by dry combustion using an elemental analyser (EuroVector, Italy; Rodrigues et al. 2009).

##### Soluble sugars content

One leaf disc (Ø 12 mm) was stored at −80 °C and used, later, to quantify the amount of soluble sugars. These were extracted from the leaf disc using 80% ethanol at 80 °C and then quantified through changes in absorbance at 405 nm, for sucrose (de Carvalho et al. 2015), and 490 nm for glucose and fructose (Santos et al. 2017).

#### Spider-mite performance

Six leaf disc arenas (Ø 12 mm) were made from one leaf of each plant (3^rd^ from below). One female spider mite of one of the two species was placed on each arena (3 arenas per species) and allowed to feed and oviposit for 4 days. Daily survival and fecundity of each female were recorded. The daily fecundity of spider mites was obtained dividing the number of eggs laid per the number of days the female lived. In a previous study, it has been shown that this measurement is highly correlated with total lifetime fecundity (Clemente et al. 2018). Therefore, this measure can also be considered as an indication of the overall performance of spider mites.

#### Interaction between Cd accumulation and organic defences

To test whether the effect of Cd and organic defences on herbivores are independent, tomato plants were exposed to 3 different Cd concentrations (0, 0.5 and 1.5 mM) as described before. Next, plants from the three treatments were infested for 48 hours with either 100 *T. evansi* or *T. urticae* females on the 3^rd^ leaf (from below), or they were left un-infested (N= 12 plants per treatment; 9 treatments: 3 Cd concentrations vs 3 infestation status). Afterwards, the plants were cleaned by removing all the mites, web and eggs with a brush.

##### The performance of spider mites

was determined as above.

##### Activity of trypsin inhibitors (TIs)

Plant material from the leaf used to determine the performance of spider mites was stored at −80 °C and used, later, to quantify the activity of TIs, as a proxy for the production of organic defences against spider mites (Sarmento et al. 2011). Approximately 300 mg of the leaf material stored at −80 °C was weighted, grounded and homogenized with 600 μL of extraction buffer (0.1M Tris-HCl, pH 8.2; 20 mM CaCl2; 1:3). Each sample was centrifuged at 4°C, 16.0xG for 25 minutes, and the supernatant was separated from the pellet and used in the spectrophotometer assay. This assay, adapted by Paulo et al. (2018) consisted in measuring the changes in absorbance at 405 nm caused by the activity of trypsin upon its substrate N-Benzoyl-D,L-arginin-4nitroamilide hydrochloride (BApNA).

### Statistical analyses

All statistical analyses were performed with the software package R 3.0.2. The normality of the residuals of each model was tested using a Shapiro-Wilk normality test and, when needed, a box-cox transformation to the data was performed. Models were simplified by sequentially removing non-significant interactions and factors. Due to logistic constrains, each experiment was repeated in blocks of 3 plants per treatment. As so, the identity of the block was included in the models as a random factor.

The effect of Cd exposure on NDVI, SC index (R1110/R810) and UV-B was determined using a general linear mixed model (lmm) with, respectively NDVI, SC or UV-B as response values, Cd supplied as a fixed factor and block as a random factor.

The relation between the Cd contained in the solution administrated to the soil and the Cd contained in the leaves was determined with a Spearman correlation, due to the non-normality of the data. Furthermore, the relation between Cd contents and the amount of calcium (Ca) and magnesium (Mg) present on the leaves was assessed with a Pearson correlation.

The effect of Cd on specific leaf area, water and soluble sugar contents was tested using a general linear mixed model (lmm) with Cd supplied as a fixed factor and block as a random factor, whereas differences in root/shoot and in C/N were determined using a generalized linear mixed model (glmm) with a binomial distribution.

The effect of Cd on daily fecundity of spider mites was determined for each range, using a general linear mixed model (lmm) with species tested and Cd supplied as fixed factors and block as a random factor. Additionally, because the soluble sugar contents and the spectral index R1110/R810 were affected by Cd, we tested whether changes in those traits influenced the daily fecundity of spider mites using a multivariate analysis of variance with distance matrices (adonis function, vegan package; Oksanen et al 2015). The fecundity of *T. evansi* and *T. urticae* were used as response factors, the amount of sucrose and glucose plus fructose or the spectral index R1110/R810, were used as the explanatory factors.

The statistical analysis of the interactions between Cd accumulation and organic defences were performed using a general linear mixed model (lmm) with daily fecundity of *T. evansi* or the amount of trypsin inhibited as response factors, Cd supplied (0mM; 0.5mM; 1.5mM) and infestation status (clean plants; plants previously infested with *T. urticae*; plants previously infested with *T. evansi*) used as fixed factors and block as a random factor.

## Results

### Effect of Cd on the performance of tomato plants

Cadmium exposure had no effect on NDVI (Table 1). However, significant differences were detected on the SC index R1100/R810, (Table 1) for plants exposed to 2 mM or 10 mM of Cd (Table 2), suggesting structural changes in the leaf cells. The same pattern was detected when analysing the narrow range of Cd concentrations (Table 1) but only for plants exposed to 1 mM and not for plants exposed to 1.5 mM (Table 2). Additionally, the UV-B reflectance of plants was significantly affected by Cd exposure (Table 1), for concentrations higher than 1 mM on the wide range (Table 2) and higher than 0.75 mM for the narrow range (Table 2).

**Table 1.** Statistical analyses of the effect of cadmium on leaf reflectance.

| Variable of interest | data subset | exploratory variable | Df | F | P value |
| --- | --- | --- | --- | --- | --- |
| NDVI | Wide range | Cd supplied | 6 | 0.21 | 0.97 |
|  | Narrow range |  | 5 | 0.83 | 0.53 |
| R1110/R810 | Wide range | Cd supplied | 6 | 3.19 | **0.015** |
|  | Narrow range |  | 5 | 4.49 | **0.001** |
| UVB | Wide range | Cd supplied | 6 | 11.44 | **<0.001** |
|  | Narrow range |  | 5 | 10.15 | **<0.001** |

**Table 2.** A posteriori contrasts on the effect of cadmium on leaf reflectance.

| Variable of interest | data subset | contrast | z value | P value |
| --- | --- | --- | --- | --- |
| R1110/R810 | Wide range | 0 mM vs 0.01 mM | 1.23 | 0.88 |
|  |  | 0 mM vs 0.1 mM | 1.80 | 0.54 |
|  |  | 0 mM vs 0.5 mM | 2.87 | 0.06 |
|  |  | 0 mM vs 1 mM | -1.89 | 0.49 |
|  |  | 0 mM vs. 2 mM | -0.18 | **0.001** |
|  |  | 0 mM vs. 10 mM | -0.13 | **0.008** |
|  | Narrow range | 0 mM vs 0.25 mM | -0.79 | 0.97 |
|  |  | 0 mM vs 0.5 mM | -0.20 | 0.99 |
|  |  | 0 mM vs 0.75 mM | 0.84 | 0.96 |
|  |  | 0 mM vs. 1 mM | -3.38 | **0.009** |
|  |  | 0 mM vs. 1.5 mM | -1.43 | 0.71 |
| UV-B | Wide range | 0 mM vs 0.01 mM | 1.67 | 0.64 |
|  |  | 0 mM vs 0.1 mM | 2.12 | 0.34 |
|  |  | 0 mM vs 0.5 mM | 3.91 | **0.002** |
|  |  | 0 mM vs. 1 mM | -0.11 | **<0.001** |
|  |  | 0 mM vs. 2 mM | -0.13 | **<0.001** |
|  |  | 0 mM vs. 10 mM | -0.17 | **<0.001** |
|  | Narrow range | 0 mM vs 0.25 mM | 1.50 | 0.66 |
|  |  | 0 mM vs 0.5 mM | 2.35 | 0.17 |
|  |  | 0 mM vs 0.75 mM | 4.07 | **<0.001** |
|  |  | 0 mM vs. 1 mM | -5.28 | **<0.001** |
|  |  | 0 mM vs. 1.5 mM | -5.74 | **<0.001** |

The concentration of Cd accumulated on tomato leaves correlated positively with the Cd concentrations that plants were exposed to, in a linear way (y = 52.299x + 21.165, rho = 0.945, P < 0.001; Fig.1). The amount of Ca and Mg on the leaves did not change significantly with Cd accumulation (R^2^ = 0.25, P = 0.11 for Ca and R^2^ = 0.23, P = 0.14 for Mg).

**Figure 1.**
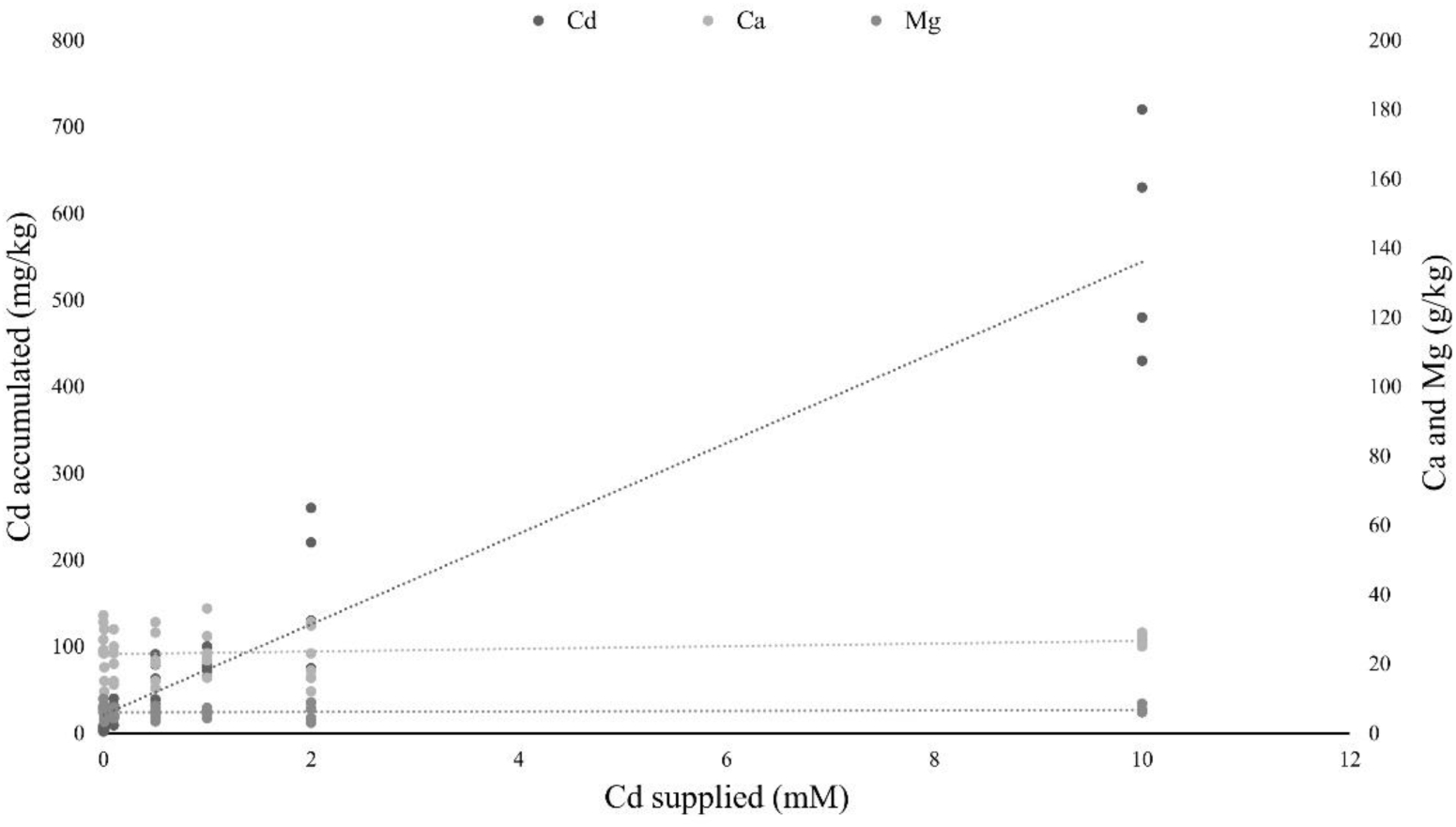
Relation between cadmium supplied in soil solution in relation to cadmium, calcium and magnesium concentration on tomato leaves. Lines represent linear regressions between the concentration of cadmium solution supplied to the plant (0, 0.01, 0.1, 0.5, 1, 2 and 10 mM; 6 plants per concentration) and the cadmium accumulated on the leaves, or the calcium and magnesium present on those leaves.

Cadmium supplied to plants did not significantly affect water content of the leaves, SLA and root/shoot (Table 3,4; Fig. 2a,b). No effect of Cd exposure was observed on the C/N content of the tomato leaves up to 1.5 mM (Table 3; Fig. 2c). However, the amount of soluble sugars on the leaves was affected by the concentration of Cd to which plants were exposed (Table 3; Fig. 2d). The amount of both sucrose and glucose plus fructose decreased in the plants exposed to low concentrations of Cd, having the lower values at 0.5 mM (Table 5; Fig. 2d). On plants exposed to 0.75 mM of Cd, the levels of sugars peeked to values higher than on control un-exposed plants but then decreased again for higher concentrations to values lower than on control plants (Table 5; Fig. 2d).

**Table 3.** Statistical analyses of the effect of cadmium on plant performance traits.

| Variable of interest | explanatory variable | Df | $\chi^2$ / F | P value |
| --- | --- | --- | --- | --- |
| Water content | Cd supplied | 5 | 34.03 | 0.31 |
| SLA (specific leaf area) | Cd supplied | 5 | 1.70 | 0.16 |
| Root/shoot | Cd supplied | 5 | 34.01 | 0.81 |
| C/N | Cd supplied | 6 | 4.42 | 0.65 |
| Sucrose | Cd supplied | 6 | 9.77 | **<0.001** |
| Glucose + fructose | Cd supplied | 6 | 13.21 | **<0.001** |

**Table 4.** Effect of cadmium on plant biomass. Average biomass of plants exposed to the narrow range of cadmium (N = 6)

| **Biomass** | **Contrast** | **Fresh weight (mg)** | **Dry weight (mg)** |
| --- | --- | --- | --- |
| Shoots | 0 mM | 48.44 ± 0.35 | 5.34 ± 0.06 |
|  | 0.1 mM | 52.23 ± 0.61 | 5.23 ± 0.09 |
|  | 0.25 mM | 41.52 ± 0.68 | 4.48 ± 0.09 |
|  | 0.5 mM | 47.00 ± 0.48 | 5.23 ± 0.08 |
|  | 0.75 mM | 37.04 ± 0.76 | 3.65 ± 0.09 |
|  | 1.5 mM | 44.83 ± 0.54 | 4.41 ± 0.08 |
| Roots | 0 mM | 91.34 ± 0.94 | 5.56 ± 0.07 |
|  | 0.1 mM | 79.23 ± 0.90 | 5.00 ± 0.08 |
|  | 0.25 mM | 88.87 ± 0.89 | 5.79 ± 0.08 |
|  | 0.5 mM | 100.41 ± 1.04 | 5.00 ± 0.09 |
|  | 0.75 mM | 78.48 ± 0.62 | 4.73 ± 0.13 |
|  | 1.5 mM | 72.98 ± 0.80 | 4.36 ± 0.08 |

**Table 5.**
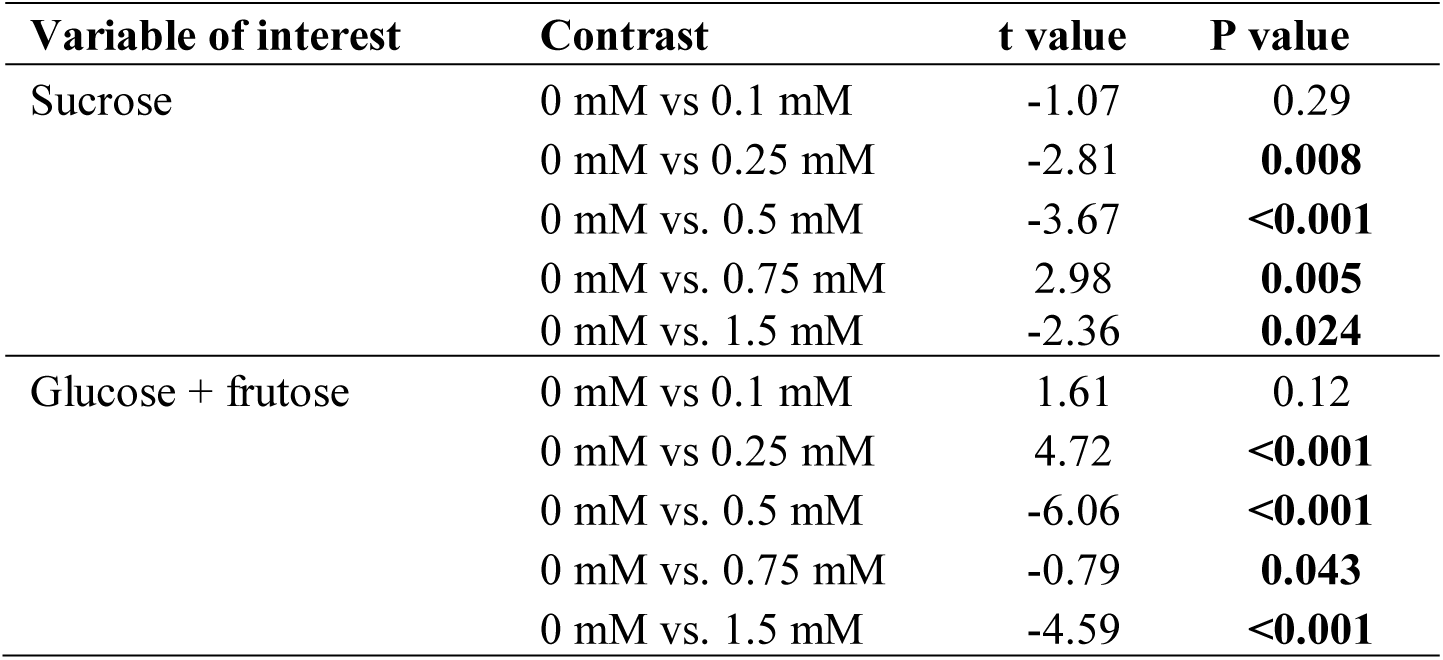
A posteriori contrasts for the effect of cadmium on soluble sugar content.

**Figure 2.**
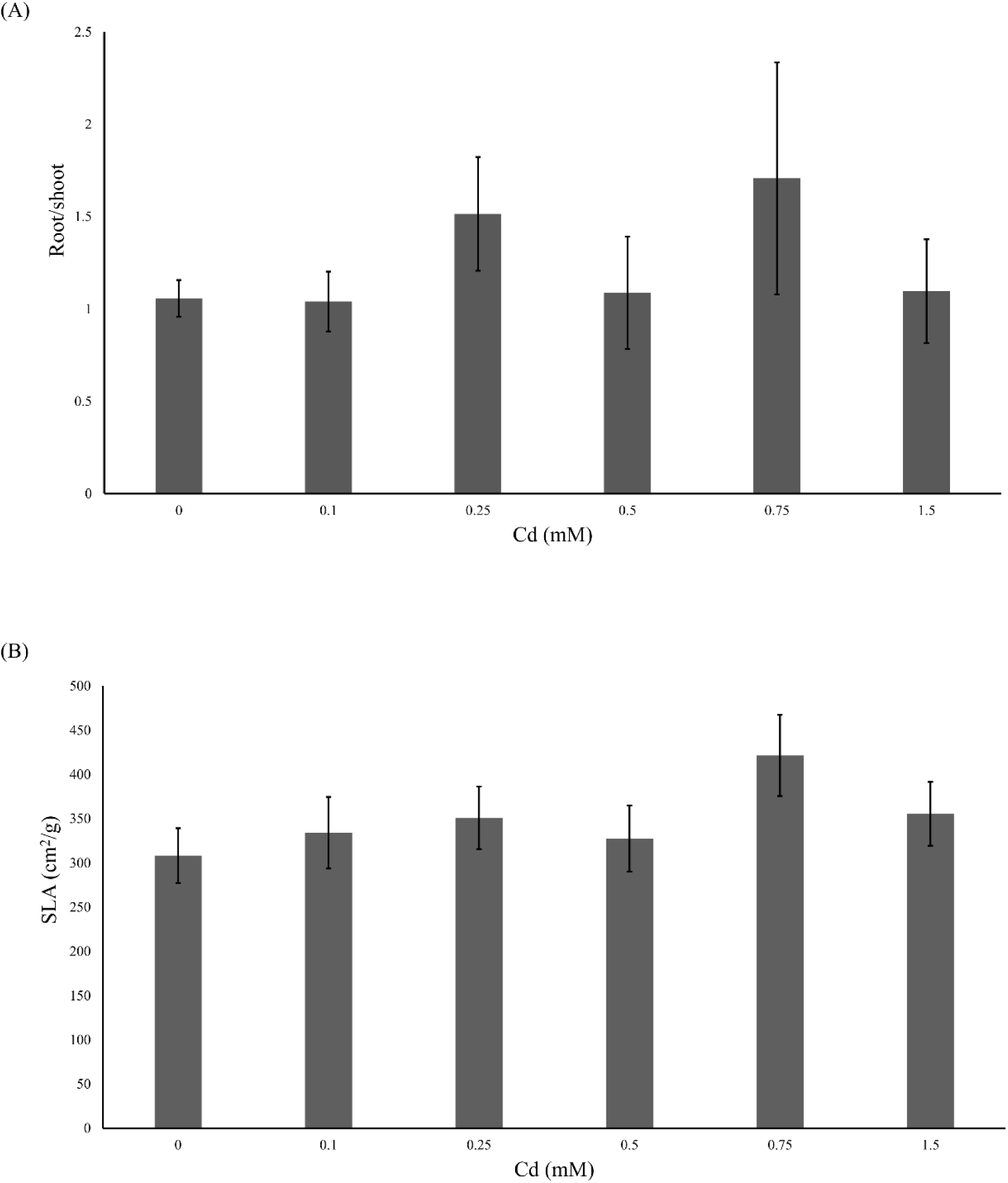
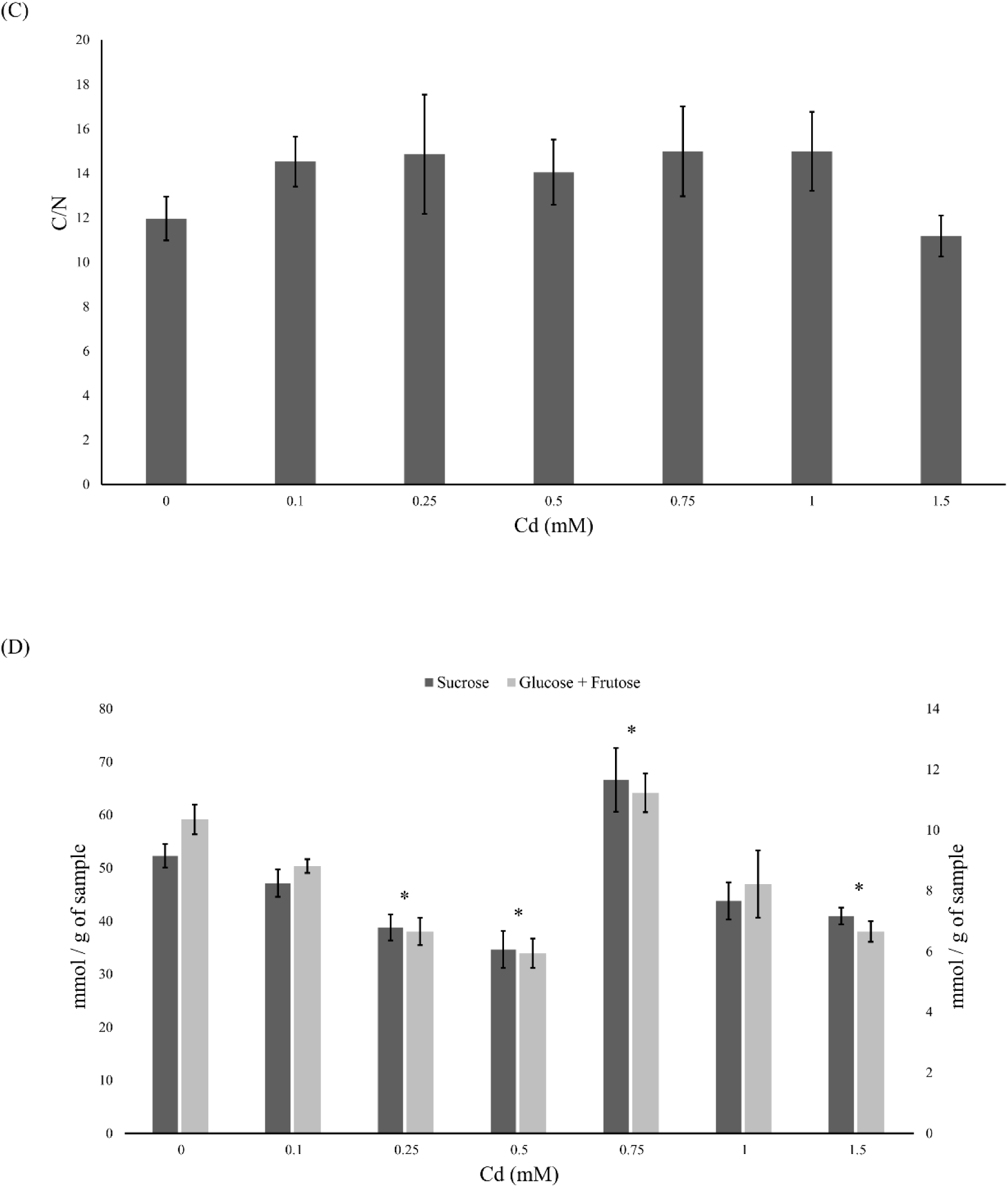
Effect of cadmium on the performance of tomato plants. Tomato plants were supplied with different cadmium concentrations (0, 0.1, 0.25, 0.5, 0.75, 1 or 1.5 mM). Plant growth (± Standard error – vertical bars; 6 plants) was measured through: a) average root to shoot ratio; b) average specific leaf area (SLA, cm^2^/g); c) average carbon to nitrogen ratio on leaves; d) average of glucose and fructose (light grey bars) and sucrose (dark grey bars) concentration (mmol per gram of fresh weight of plant). ^*^ represent significant differences from the control plants.

### Effect of Cd accumulation on the performance of spider mites

The oviposition of spider mites on leaf discs was significantly affected by the Cd supplied to the plants used to make those discs (Table 6, Fig.3). Additionally, both spider-mite species were similarly affected by the Cd concentration that plants were exposed to (Table 6). Both species increased their oviposition with low amounts of Cd until a threshold concentration, 0.5 mM (Table 6, Fig. 3). From this concentration onwards, Cd had a negative effect on the oviposition rate of spider mites, reaching, for the wide range, values lower than on control plants at 2 mM and on the narrow range at 1.5mM (Table 6; Fig.3).

**Table 6.** Statistical analyses on the effect of cadmium on the performance of spider mites.

| Variable of interest | data subset | explanatory variable | Df | F | P value |
| --- | --- | --- | --- | --- | --- |
| Daily fecundity | Wide range | Cd supplied x tested species | 6 | 47.04 | 0.14 |
|  |  | Cd supplied | 6 | 64.13 | **<0.001** |
| Daily fecundity | Narrow range | Cd supplied x tested species | 6 | 0.12 | 0.99 |
|  |  | Cd supplied | 6 | 10.62 | **<0.001** |
| Variable of interest | data subset | contrast |  | Z value | P value |
| Daily fecundity | Wide range | 0 mM vs 0.01 mM | - | 0.32 | 0.99 |
|  |  | 0 mM vs 0.1 mM |  | -0.94 | 0.96 |
|  |  | 0 mM vs. 0.5 mM | - | -3.14 | **0.028** |
|  |  | 0 mM vs. 1 mM |  | -1.34 | 0.83 |
|  |  | 0 mM vs. 2 mM | - | -3.99 | **0.012** |
|  |  | 0 mM vs. 10 mM | - | -4.82 | **<0.001** |
| Daily fecundity | Narrow range | 0 mM vs 0.1 mM | - | -0.58 | 0.99 |
|  |  | 0 mM vs 0.25 mM | - | -1.45 | 0.77 |
|  |  | 0 mM vs. 0.5 mM | - | -3.26 | **0.018** |
|  |  | 0 mM vs 0.75 mM | - | 0.50 | 0.99 |
|  |  | 0 mM vs. 1 mM | - | -1.43 | 0.78 |
|  |  | 0 mM vs. 1.5 mM | - | 3.93 | **0.001** |

**Figure 3.**
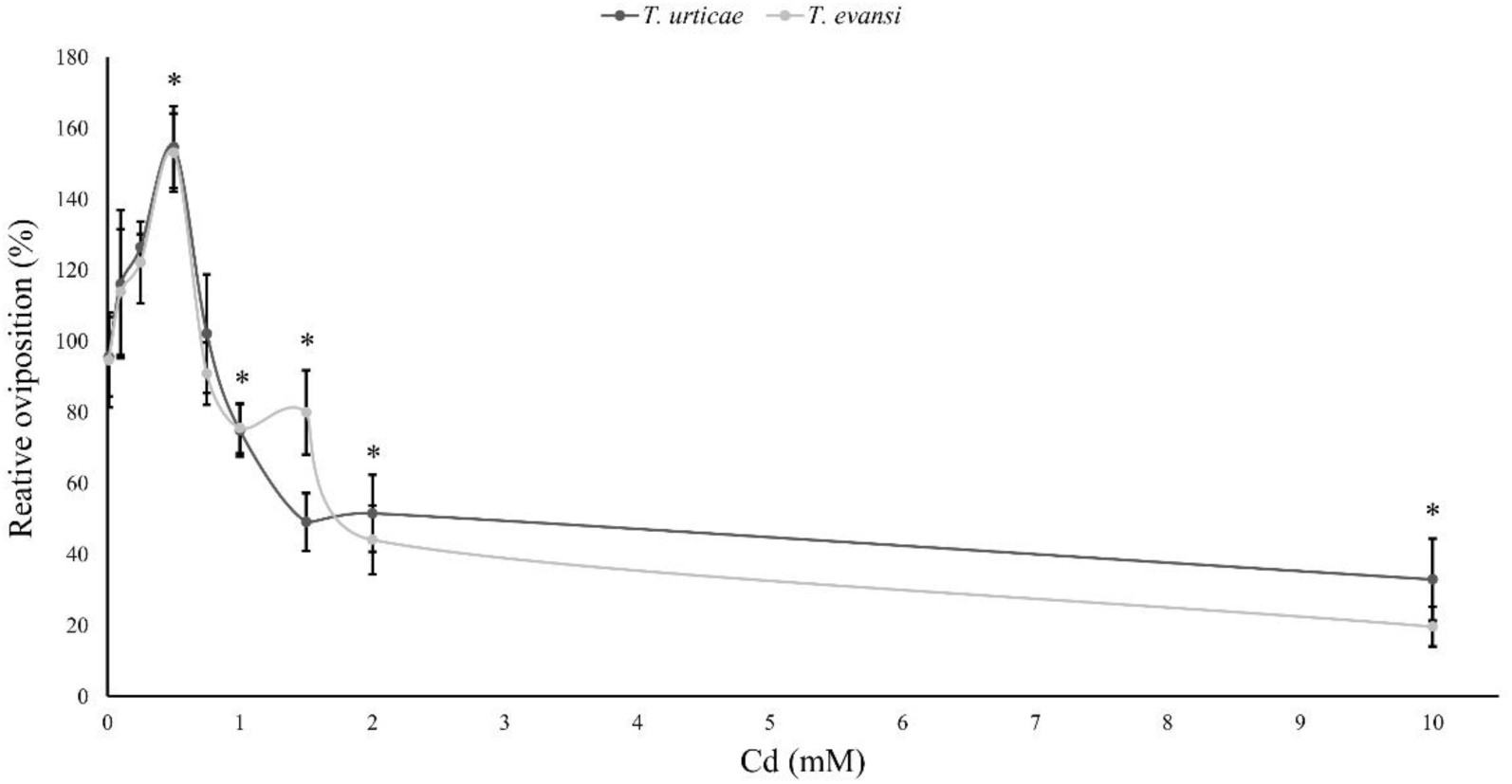
Performance of spider mites on leaves of tomato exposed to cadmium. Average relative oviposition rate of *T. evansi* (light grey) and *T. urticae* (dark grey) females on tomato plants (± Standard error – vertical bars; 6/12 plants, 3 discs per species per plant). For each range of cadmium solutions, a (0, 0.01, 0.1, 0.5, 1, 2 or 10 mM, N= 6) and b (0, 0.1, 0.25, 0.5, 0.75, 1 or 1.5 mM, N= 12) the oviposition of spider mites was normalized to the control (no cadmium) and merged in the same panel.

The amount of sucrose on the leaves did not affect the fecundity of spider mites (F_1_ = 0.008, P = 0.71). In contrast, the amount of glucose plus fructose affected this trait (F_1_ = 0.19, P = 0.003). Additionally, the reflectance index R1110/R810 affected the fecundity of spider mites (F_1_ = 0.07, P = 0.005).

### Interaction between Cd accumulation and organic defences

The oviposition rate of *T. evansi* was affected by both Cd concentration and previous infestation with conspecifics or heterospecifics (Table 7; Fig. 4). However, the interaction between these effects was not significant (Table 7). The oviposition rate of *T. evansi* increased with previous infestation by conspecifics and decreased with previous infestation by *T. urticae* (Table 7; Fig. 4), independently of the concentration of Cd to which plants were exposed before. Moreover, the oviposition rate of *T. evansi* increased on plants exposed to 0.5 mM of Cd and decreased on plants exposed to 1.5 mM of Cd (Table 7; Fig. 4), compared to control plants, as observed in the previous results (Fig. 3).

**Table 7.** Statistical analyses of the effect of cadmium and spider mite infestation.

| Variable of interest | explanatory variable | Df | F | P value |
| --- | --- | --- | --- | --- |
| Daily fecundity | Cd supplied x infestation status | 4 | 0.47 | 0.76 |
|  | Infestation status | 2 | 27.56 | **<0.001** |
|  | Cd supplied | 2 | 32.18 | **<0.001** |
| Trypsin inhibitors | Cd supplied x infestation status | 4 | 0.03 | 0.97 |
|  | Infestation status | 2 | 4.49 | **0.014** |
|  | Cd supplied | 2 | 0.77 | 0.38 |
| Variable of interest | contrast |  | z value | P value |
| Daily fecundity | Un-infested vs. *T. evansi* | - | -3.22 | **0.021** |
|  | Un-infested vs. *T. urticae* | - | 3.49 | **0.009** |
|  | 0 mM vs. 0.5 mM | - | 4.19 | **<0.001** |
|  | 0 mM vs. 1.5 mM | - | -3.88 | **<0.001** |
| Trypsin inhibitors | Un-infested vs. *T. evansi* | - | -3.26 | **0.018** |
|  | Un-infested vs. *T. urticae* | - | 3.93 | **0.021** |

**Figure 4.**
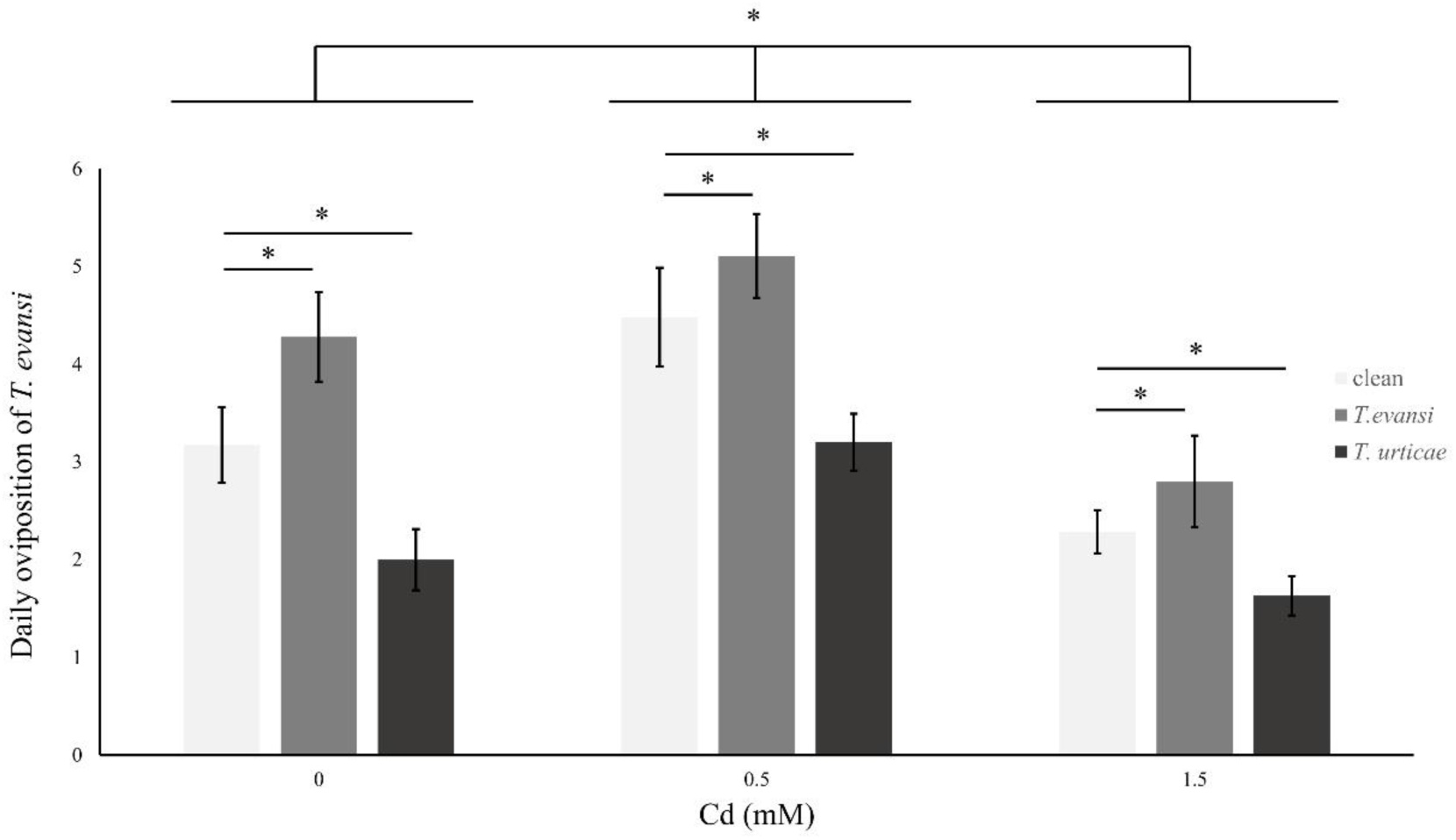
Effect of cadmium exposure and herbivory on the performance of subsequent infestations. Average number of eggs laid per day *T. evansi* females on clean plants (light grey), plants infested with 100 *T. evansi* females (grey) or with 100 *T. urticae* females (dark grey) for 48 hours. Plants (± Standard error – vertical bars; 12 plants, 3 discs per species per plant) were exposed to a range of cadmium concentrations (0, 0.5 or 1.5 mM). ^*^ represent significant differences between infestation treatments and between cadmium treatments, there was a non-significant interaction between the two factors.

Additionally, the activity of trypsin inhibitors was affected by infestation by spider mites independently of the concentration of Cd exposed to the plants (Table 7, Fig. 5). Cadmium accumulation had no significant effect on the activity of trypsin inhibitors (Table 7; Fig. 5).

**Figure 5.**
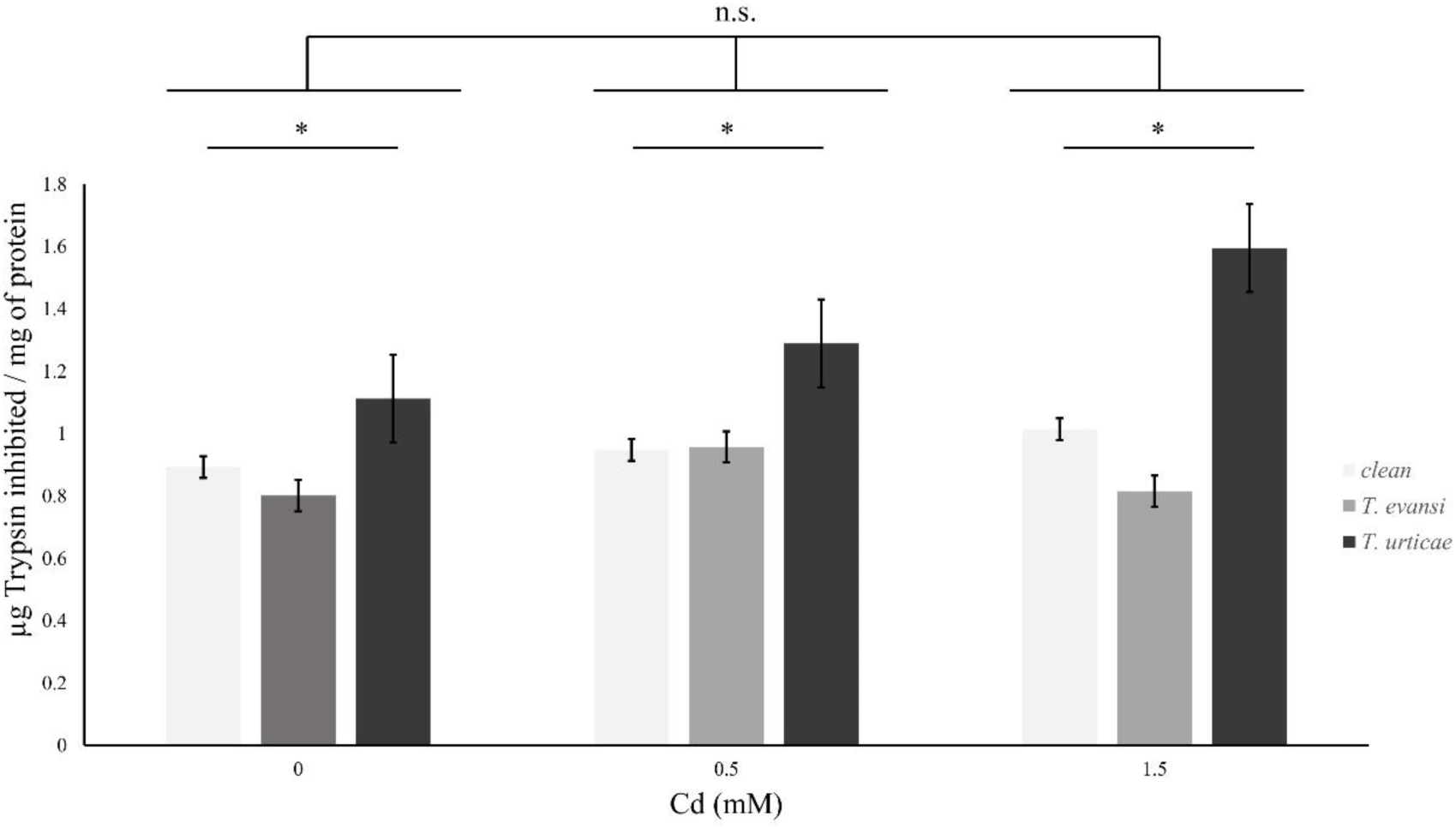
Effect of cadmium exposure and herbivory on organic plant defences. Amount (µg) of trypsin inhibited per mg of protein in leaf samples of clean plants (light grey), plants infested by 100 *T. evansi* (grey) females for 48 hours or 100 *T. urticae* (dark grey) females for 48 hours. Plants (± Standard error – vertical bars; 12 plants, 3 discs per species per plant) were exposed to a range of Cd concentrations (0, 0.5 or 1.5 mM). ^*^ represent significant differences between infestation treatments, Cd supplied had no significant effect, neither did was significant the interaction between the two factors.

## Discussion

Within the tested ranges Cd exposure did not affect tomato specific leaf area, root/shoot, water content (Fig. 2) or NDVI, revealing no effect on plant growth. However, the soluble sugar content was affected by Cd exposure, as was the reflectance index R1110/R810, suggesting structural changes on leaf cells. Spider mites infesting those plants were affected by Cd concentrations, albeit in a non-linear way (Fig. 3). Indeed, both spider mite species had increased performance on plants mildly exposed to Cd, as compared to un-exposed plants, but lower performance after a given threshold, revealing a hormetic effect. Finally, the interaction of both spider mites with plant defences was not affected by the level of Cd that the plants were exposed to (Fig. 5). Together, these results suggest that metal accumulation and the production of plant defences against herbivores do not interact with each other.

Studies regarding Cd accumulation by tomato plants reveal high variability in this trait (Hartke et al. 2013), with some plants accumulating amounts below the hyperaccumulation threshold (<100 mg/kg, Pollard 2000), even at high concentrations of Cd supply (Ammar et al. 2007, 2008) and others accumulating above this threshold (Dong et al. 2006, Gratão et al. 2008, López-Millán et al. 2009). Here we observe that Cd accumulated linearly in the leaves of tomato plants (Fig. 1), up to values above the hyperaccumulation threshold, suggesting that, *Moneymaker*, the variety of tomato used in our study, is as a facultative hyperaccumulator. This is further confirmed by the values of Ca and Mg on the leaves which remain stable with increasing Cd in the leaves, as seen for other hyperaccumulator plants (Gomes et al. 2013, Pereira et al. 2017).

Here we report the absence of an immediate negative impact on plant growth. Moreover, we also observe no differences in the carbon to nitrogen ratio on leaves of plants exposed to different Cd concentrations, indicating no shifts in the growth/defence balance (Herms and Mattson 1992). This contrasts with previous studies showing a negative impact of Cd on tomato plant growth, for Cd accumulation values within the ranges used here (Dong et al. 2006, Ammar et al. 2007, Gratão et al. 2008, López-Millán et al. 2009). Possibly, the variety of tomato we used in this experiment is more tolerant to Cd than most other varieties. Indeed, the few studies using this variety observe no signs of toxicity (Petit and Geijn 1978, Petit et al. 1978). Another possibility is that the growing substrate affected these results. Indeed, most studies on this topic used continuously aerated hydroponics, creating an artificial situation for the plants, and here we used soil as a substrate as in natural conditions. Growing in soils may be advantageous to plants, given that soil microbiota may regulate the process of metal accumulation in the shoots (de Souza et al. 1999, Farinati et al. 2009), reducing the costs involved in this process for the plant (Farinati et al. 2009).

In contrast to most plant traits that did not respond to Cd, we found changes in soluble sugar contents and leaf reflectance. Changes in the amount of soluble sugars in the leaves with Cd were non-linear. Soluble sugars are generally associated with an initial response to plant stress, with changes in their accumulation, either increasing or decreasing, affecting the REDOX reactions originated by environmental stress (Couée et al. 2006). Cadmium supply may lead to either an increase (Mishra et al. 2014) or a decrease (Scheirs et al. 2016, Shackira and Puthur 2017) in the amount of soluble sugars in the shoots of exposed plants. The fluctuations we observe in the soluble sugars content may indicate that these are being affected by different processes in the plant, and this may help to reconcile the contrasting observations in the literature. Indeed, plants may also have an hormetic response to abiotic stressors, increasing their performance with small amounts of Cd until a threshold where the negative effects caused by this metal are higher than the positive (Siddhu et al. 2008). Corroborating this hypothesis requires more controlled experiments and a systematic measurement of Cd concentrations in the leaves. Moreover, higher concentrations of Cd exposure caused changes in the leaf reflectance (R1110/R810), which have been linked to structural changes on their leaf cells (Sridhar et al. 2007). Additionally, plants exposed to the higher concentrations of Cd showed a significant increase in the reflectance of UV-B light, which is possibly related to the production of phenolic compounds that protect and signal plants against abiotic stresses (Roberts and Paul 2006, Izaguirre et al. 2007). Our results thus highlight the need to collect different measures of plant performance for a single abiotic stress.

The performance of spider mites was also affected by Cd accumulation in tomato leaves. Both species had a non-linear, hormetic response to this metal. Most herbivores are negatively affected by metal accumulation in the leaves of their host plants (Hanson et al. 2003, Freeman et al. 2007, Quinn et al. 2010, Stolpe et al. 2017). Still, there are some examples of higher abundances of herbivores on sites with intermediate concentrations of toxic metals (Zvereva et al. 1995, Kozlov et al. 2003), under natural conditions. Nevertheless, this is, to our knowledge, the first report of an hormetic effect on the herbivores performance with increasing metal concentrations in the plant, observed in controlled conditions. This phenomenon may have an important role in the evolution of metal accumulation, as it may pose a strong selective pressure to the plants to accumulate higher amounts of metal and surpass the beneficial effect of mild doses to the herbivores.

This hormetic pattern may be due to direct effects of the metal on the spider mites, or indirectly, through changes in plant quality. We observed no effect of Cd on plant biomass and C/N ratio and, thus, the growth/defence balance was not affected. However, the amount of soluble sugars in the leaf significantly affected the performance of spider mites, revealing indirect effects of plant quality. Yet, the performance of spider mites has been reported as positively (Ximénez-Embún et al. 2016, 2017) or negatively (Wermelinger et al. 1985, Joutei et al. 2000, Scheirs et al. 2006) correlated to sugar content, indicating that this may be dependent on the host plant species or on other physiological responses that were not assessed. Additionally, we observed that the daily fecundity of both species correlated with the leaf spectral index R1110/R810, indicating a possible effect of structural changes on leaf cells caused by Cd. Still, our experiments do not exclude a possible direct effect of the metal on the performance of spider mites. Whether these correlations imply causality is a relevant question that calls for future studies.

The similarity in the hormetic pattern of the two species suggests that they have overlapping fundamental niches, potentially increasing the competition between them. Indeed, if mites perform better at intermediate Cd concentrations, they may prefer to establish on those plants rather than on un-contaminated plants. Moreover, their higher oviposition on those plants is probably indicative of a higher growth rate (Clemente et al. 2018). This may entail a faster saturation of that environment, relative to others. If this is the case, then plants are expected to pay a high cost of mild Cd accumulation. Thus, given enough time, plants may be selected to ‘avoid’ the level of Cd accumulation that results in better performance for the herbivores, being selected to accumulate higher amounts of metal, becoming hyperaccumulators, or to not accumulate metal at all, becoming excluders.

Because the two spider mites have dissimilar interactions with tomato organic defences, the fact that they perform best on plants with the same Cd supply also suggests no interaction between metal accumulation and the production of plant defences. Furthermore, the contrasting effects of these two spider-mite species on the activity of trypsin inhibitors and its effect on subsequent infestations were consistent across Cd environments, revealing no interference of Cd on protease activity, in contrast to what was seen in other plant species (Pena et al. 2006, Lin et al. 2010). The plants used in these experiments showed little evidence of Cd toxicity. Possibly, the effect of Cd was not strong enough to induce the protective protease activity reported for other plants (Pena et al. 2006, Lin et al. 2010). Additionally, the effect of metal supply on spider-mite performance was not affected by previous infestation. Together, these results suggest that metal-based and organic plant defences do not interfere with each other, serving the same purpose. Few studies have addressed this potential interference. Some studies reveal that the expression of organic defences is lower with high metal supply (Davis et al 2001, Tolra et al. 2001, Farinati et al. 2009, 2011, Fones et al. 2013). However, these studies concern plants that are obligate hyperaccumulators, which occur exclusively in contaminated soils. If metal accumulation provides the same function as organic defences, and if the production of organic defences is costly, this may select for a reduction in organic defences in plants under high metal supply. The opportunity for such selection to be effective is much higher on obligate metal accumulators (Poschenrieder et al. 2006), which is not the case of tomato. Alternatively, plants may be suffering from metal accumulation, hence they may lack the necessary resources to trigger organic defences (Farinati et al. 2009, 2011, Fones et al. 2013). As we did not observe negative effects of Cd on plant growth, it may be that cost on tomato plants due to metals was not sufficient to lead to a trade-off between these two types of defences. Possibly, long-term exposure to this contaminant, or exposure to higher concentrations, would cause significant costs to the plant affecting its growth rate or posing constrains in fruit production. Still, a recent field work found little evidence for these trade-offs (Kazemi-dinan et al. 2015).

Another possible explanation for the absence of a trade-off in our study is that the effect of metal accumulation on herbivores was non-linear. Thus, if plants would produce fewer organic plant defences, as metal accumulation increased, herbivores would have an extra-advantage at intermediate metal concentrations, benefiting from both a high performance in response to metals, and a low exposure to organic defences. This, in turn, would pose a strong selective pressure upon plants not to shut down organic defences. In the absence of an interaction between metal-based and organic defences, plants occurring in heterogeneous environments may fine tune these strategies depending on their relative performance on each environment. Possibly, plants accumulate more metal when exposed to herbivores that suppress their organic defences, overcoming the positive effects that low concentrations may have on these herbivores. This hypothesis awaits to be tested.

In sum, our results suggest that the community of spider mites on tomato plants will be similar on contaminated and un-contaminated soils. Yet, whether spider mites with different interactions with organic plant defences adapt at a similar rate to plants with metals remains an open question. Our results shed some light on the interactions between plants and herbivores on metal polluted environments. More studies are now needed to fully grasp the relationship between metal accumulation and the production of organic defences on plants that are facultative hyperaccumulators.

## Acknowledgements

We would like to thank Plant Biology Department of the Sciences Faculty of the University of Lisbon for the space we used in the plant climatic chamber, Inês Santos for helping with plant growing and maintenance of spider mite populations, Alice Nunes for support in the statistical analyses, and all members of “Adaptation to heterogenous environments” and “eChanges” groups for stimulating discussions and scientific inputs. Funds were provided by an ERC consolidator grant (COMPCON, GA 725419) to SM, by UID/BIA/00329/2013 (2015-2017) and UID/MULTI/04046/2013 center grants from FCT, Portugal to cE3c and BioISI respectively, and by an FCT PhD scholarship PD/BD/114010/2015 to DG.

The authors declare no conflict of interest. All data used in this work will be archived in Dryad upon acceptance of the manuscript for publication.

DG, SM and CB conceived the study. DG, SM and CB designed the experiments with help from HCS and AS. DG collected the data with assistance from HCS and AS. DG analysed the data with help from all authors. DPG and SM led the writing of the manuscript, with significant help from CB and contributions by all authors. All authors gave final approval for publication.

